# Effect of channel assembly (KCNQ1 or KCNQ1 + KCNE1) on the response of zebrafish I_Ks_ to I_Ks_ inhibitors and activators

**DOI:** 10.1101/2021.10.26.465845

**Authors:** Jaakko Haverinen, Minna Hassinen, Matti Vornanen

## Abstract

In cardiac myocytes, the slow component of the delayed rectifier K^+^ current (I_Ks_) ensures repolarization of action potential during beta-adrenergic activation or when other repolarizing K^+^ currents fail. As a key factor of cardiac repolarization I_Ks_ should be present in model species used for cardiovascular drug screening, preferably with pharmacological characteristics similar to those of the human I_Ks_. To this end, we investigated the effects of inhibitors and activators of the I_Ks_ on KCNQ1 and KCNQ1+KCNE1 channels of the zebrafish, an important model species, in Chinese hamster ovary cells. Inhibitors of I_Ks_, chromanol 293B and HMR-1556, inhibited zebrafish I_Ks_ channels with approximately similar potency as that of mammalian I_Ks_. Chromanol 293B concentration for half-maximal inhibition (IC_50_) of zebrafish I_Ks_ was at 13.1±5.8 and 13.4±2.8 μM for KCNQ1 and KCNQ1+KCNE1 channels, respectively. HMR-1556 was a more potent inhibitor of zebrafish I_Ks_ with IC_50_=0.1±0.1 μM and 1.5±0.8 μM for KCNQ1 and KCNQ1+KCNE1 channels, respectively. R-L3 and mefenamic acid, generally identified as I_Ks_ activators, both inhibited zebrafish I_Ks_. R-L3 almost completely inhibited zebrafish I_Ks_ generated by KCNQ1 and KCNQ1+KCNE1 channels with similar affinity (IC_50_ 1.1±0.4 and 1.0±0.4 μM, respectively). Mefenamic acid partially blocked zebrafish KCNQ1 (IC_50_=9.5±4.8 μM) and completely blocked KCNQ1+KCNE1 channels (IC_50_=3.3±1.8 μM). Although zebrafish I_Ks_ responds to I_Ks_ inhibitors in the same way as mammalian IKs, its response to activators is atypical, probably due to the differences in the binding domain of KCNE1 to KCNQ1. Therefore, care must be taken when translating the results from zebrafish to humans.

## INTRODUCTION

The zebrafish (*Danio rerio*) is an established animal model in developmental biology, genetics and several other disciplines because of its research technical, economical, and ethical benefits (Kari et al., 2007; Verkerk and Remme, 2012; Vornanen et al., 2018; Narumanchi et al., 2021). For the same reasons, zebrafish has been sometimes considered as the optimal animal model for preclinical drug screening (Parng et al., 2002; MacRae and Peterson, 2015; de la Cruz et al., 2020). The primary goal of the preclinical drug screening “is to determine if the product is reasonably safe for initial use in humans, and if the compound exhibits pharmacological activity that justifies commercial development” (U.S. Food and Drug Administration, Investigational New Drug (IND) Application). In order to obtain significant results from preclinical studies with high generalizability, appropriate animal models that are as comparable as possible to the target human population are required (Honek, 2017). The requirements for preclinical drug screening are much more stringent than for basic science (Wall and Shani, 2008). It is not enough that humans and model species share common principles of gene function and regulation, the binding affinity of a drug and its effect on targets should also be quantitatively similar in humans and model species (Wall and Shani, 2008; Pound and Ritskes-Hoitinga, 2018). If those requirements are not met, it is possible that useful molecules - which appear toxic in model species - might be rejected from the drug development pipeline as false positives, or conversely, toxic compounds to humans, but not to the model animal, might proceed to clinical drug development program (Wall and Shani, 2008). Therefore, species selection should be based on what gives the best and safest platform for human testing (Honek, 2017). In cardiac electrophysiology, the zebrafish is in some respects a better model than the mouse. Unlike the mouse ventricular action potential (AP), the zebrafish ventricular AP has a clear plateau phase, and the heart rate of the zebrafish is more reminiscent of the human heartbeat (Brette et al., 2008; Nemtsas et al., 2010). In addition, repolarization of zebrafish and human cardiac AP is largely based on the fast (I_Kr_) and slow components (I_Ks_) of the delayed rectifier K^+^ current rather than transient outward current (I_to_) and ultra-rapid K^+^ current (I_Kur_) typical of the murine heart (Xu et al., 1999; Verkerk and Remme, 2012; Vornanen and Hassinen, 2016; Joukar, 2021). Despite those similarities in ion current composition, drug responses of the zebrafish ion channels are still poorly understood. Because the pharmacological properties of the zebrafish I_Ks_ have not been earlier examined, we decided to characterize the responses of zebrafish I_Ks_ channels to identified inhibitors and activators of the I_Ks_.

In addition to I_Kr_, I_Ks_ is the main regulator of ventricular AP duration and thus QT time in the electrocardiogram. Under normal circumstances, I_Ks_ plays little role in ventricular AP repolarization (Bendahhou et al., 2005; Bett et al., 2006). However, when ventricular repolarization faces challenging situations, notably under stressful conditions of increased sympathetic tone, I_Ks_ contribution is markedly increased at high heart rates (Rocchetti et al., 2001; Jost et al., 2005). Thus, under vulnerable conditions, I_Ks_ provides a repolarization reserve that can prevent excessive AP prolongation and development of arrhythmogenic afterdepolarizations, especially when the other repolarizing current I_Kr_ is compromised (Chen et al., 2003; Bendahhou et al., 2005; Bett et al., 2006). In mammalian hearts, I_Ks_ is generated by a channel assembly consisting of the KCNQ1 alpha subunits and KCNE1 (MinK) beta subunits, although the exact stoichiometry of the subunits is disputed (Sanguinetti et al., 1996; Barhanin et al., 1996; Chen et al., 2003; Morin and Kobertz, 2008; Wang et al., 2011). Expression of KCNQ1 alone generates a rapidly activating and inactivating delayed-rectifier K^+^ current, whose properties do not match with those of the native cardiac I_Ks_. Assembly of KCNQ1 with KCNE1 profoundly modifies the biophysical characteristics of KCNQ1 including activation and deactivation kinetics, frequency-response and beta-adrenergic responsiveness. In particular, the presence of KCNE1 in the channel assembly affects drug binding of the channel (Bett and Rasmusson, 2008).

Given that preclinical screening of cardiovascular drugs requires detailed information on the used biological targets, it is astonishing how little attention has been paid to the comparison of human and zebrafish cardiac ion currents/channels, the main targets of arrhythmia medicines. Understanding the molecular basis of subunit–channel interactions and their impact on drug responses is therefore of critical importance when model organisms are used for preclinical drug screening. While it is known that gating kinetics and pharmacological interactions of I_Ks_ channel is strongly affected by the presence of the KCNE1 beta subunit (Barhanin et al., 1996; Bett et al., 2006; Lerche et al., 2007; Bett and Rasmusson, 2008; Hassinen et al., 2011), the effect of channel assembly on drug responses in zebrafish has not been studied. We recently indicated that I_Ks_ is present is zebrafish ventricular myocytes and it affects the duration of ventricular AP (Abramochkin et al., 2018). Notably, transcripts of the KCNQ1 are expressed at much higher levels than those of KCNE1, and electrophysiological properties of the ventricular I_Ks_ suggest that many of the channels might be homotetrameric KCNQ1 channels. Therefore, we decided to examine how the subunit composition of the zebrafish I_Ks_ channel affects the currents response to identified I_Ks_ inhibitors and activators. To this end, we expressed zebrafish KCNQ1 and KCNQ1+KCNE1 channels in CHO cells and examined their responses to known activators and inhibitors of the mammalian I_Ks_. Because subunit composition of the zebrafish I_Ks_ channel may contain fewer KCNE1 subunits than the mammalian I_Ks_ channel, it was hypothesized that drug responses, in which stoichiometry of subunits is crucial, would differ between mammalian and zebrafish I_Ks_.

## MATERIALS AND METHODS

### Heterologous gene expression

Zebrafish KCNQ1 and KCNE1, previously cloned to pcDNA3.1 vector (Abramochkin et al., 2018), were expressed in Chinese hamster ovary (CHO) cells. CHO cells were grown in Ham’s F12 nutrient mixture (Sigma-Aldrich) supplemented with 10% heat-inactivated fetal bovine serum (Sigma-Aldrich) and 100 U ml^−1^ penicillin-streptomycin (Thermo Scientific) at +37°C under 7.5% CO_2_ atmosphere. For transient expression of KCNQ1 and KCNE1, cells were transfected with plasmids containing either KCNQ1 alone or KCNQ1:KCNE1 in ratio 3:1 using Turbofect transfection reagent (Thermo Scientific) (Hassinen et al., 2015; Abramochkin et al., 2018). Green fluorescent protein (GFP)-coding plasmid peGFP-N1 was used to see the transfection status of the cells.

### Whole-cell patch-clamp

Whole cell patch-clamp experiments were conducted 48–56 h after transfection. The slow component of the delayed rectifier K^+^ current (I_Ks_) was recorded at 28 °C as previously reported in detail (Hassinen et al., 2011; Abramochkin et al., 2018). Cells were superfused with a saline solution containing (in mmol l^−1^): 150 NaCl, 3 KCl, 1.8 CaCl_2_, 1.2 MgCl_2_, 10 HEPES and 10 glucose, with pH adjusted to 7.7 at 20 °C with NaOH. Patch pipettes were filled with K^+^-based electrode solution containing (in mmol l^−1^): 140 KCl, 4 MgATP, 1 MgCl_2_, 5 EGTA, 0.3 Na_2_GTP, and 10 HEPES with pH adjusted to 7.2 at 20 °C with KOH. For current-voltage dependence, I_Ks_ was elicited from the holding potential of −80 mV by 5-s depolarizing pulses to −40 to +80 mV in 20-mV steps. To generate concentration-response curves, cells were exposed to cumulatively increasing concentrations of I_Ks_ current inhibitors and activators for 5 minutes during which time a steady-state response was obtained.

### Drugs

Two I_Ks_ inhibitors, chromanol 293B (*trans*-*N*-[6-Cyano-3,4-dihydro-3-hydroxy-2,2-dimethyl-2*H*-1-benzopyran-4-yl]-*N*-methyl-ethanesulfonamide) and HMR-1556 (*N*-[(3*R*,4*S*)-3,4-Dihydro-3-hydroxy-2,2-dimethyl-6-(4,4,4-trifluorobutoxy)-2*H*-1-benzopyran-4-yl]-*N*-methylmetanesulfonamide) and two I_Ks_ activators, L-364,373 (R-L3 or 5-(2-Fluorophenyl)-1,3-dihydro-3-(1*H*-indol-3-ylmethyl)-1-methyl-2*H*-1,4-benzodiazepin-2-one) (Tocris Cookson; Bristol, UK) and mefenamic (dimethylphenylaminobenzoic) acid (Sigma-Aldrich) were used. Stock solutions of all drugs were made in DMSO at concentrations of 30 mM, 1mM, 1mM, 30mM for chromanol 293B, HMR-1556, R-L3 and mefenamic acid, respectively. Working solutions were made daily in the external saline from these stock solutions.

### Data analysis and Statistics

The results are presented as means ± SEM from n cells. For the concentration-response analysis the normalized I_Ks_ current was plotted as a function of drug concentration and fitted to the sigmoidal equation

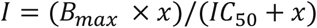

where *I* is current, *B_max_* maximum inhibition of the current, *IC_50_* concentration of the drug which causes half-maximal inhibition of the current and *x* is the drug concentration. Statistical analysis was performed with SPSS 27.0 software (IBM). After testing the normal distribution of the data, non-paired *t*-test was used to compare current densities and IC_50_ values between KCNQ1 and KCNQ1+KCNE1 channels. A *p* value < 0.05 was considered statistical significantly different.

## RESULTS

### Properties of the current generated by homomeric KCNQ1 and heteromeric KCNQ1+KCNE1 channels

There was marked differences in amplitude and kinetics of outward currents generated by homomeric KCNQ1 and heteromeric KCNQ1+KCNE1 channels. At the end of the 5-s depolarizing pulse, the density of the current produced by homomeric KCNQ1 α-subunits was about 6 times lower (23.0 ± 6.6 pA pF^−1^) than that of KCNQ1+KCNE1 heteromeric channels (142 ± 37.7 pA pF^−1^) (*p*<0.05) (**Figure 1A-C**). In addition, the activation kinetics of KCNQ1 channels were much faster with a peak current at 18.1 ± 1.8 ms, followed by a slower inactivation to the steady-state level. KCNQ1+KCNE1 channels activated much more slowly with an activation time constant (τ) of 33.7 ± 7.3 ms and showed no inactivation during depolarization. The density of the early peak of KCNQ1 current was 37.3 ± 6.7 pA pF^−1^, which is 62% larger than the steady-state current at the end of the depolarizing pulse. Cells transfected with GFP plastids had virtually no current when perfused with vehicle-only saline (DMSO) (**Figure 1C**).

**FIGURE 1.**
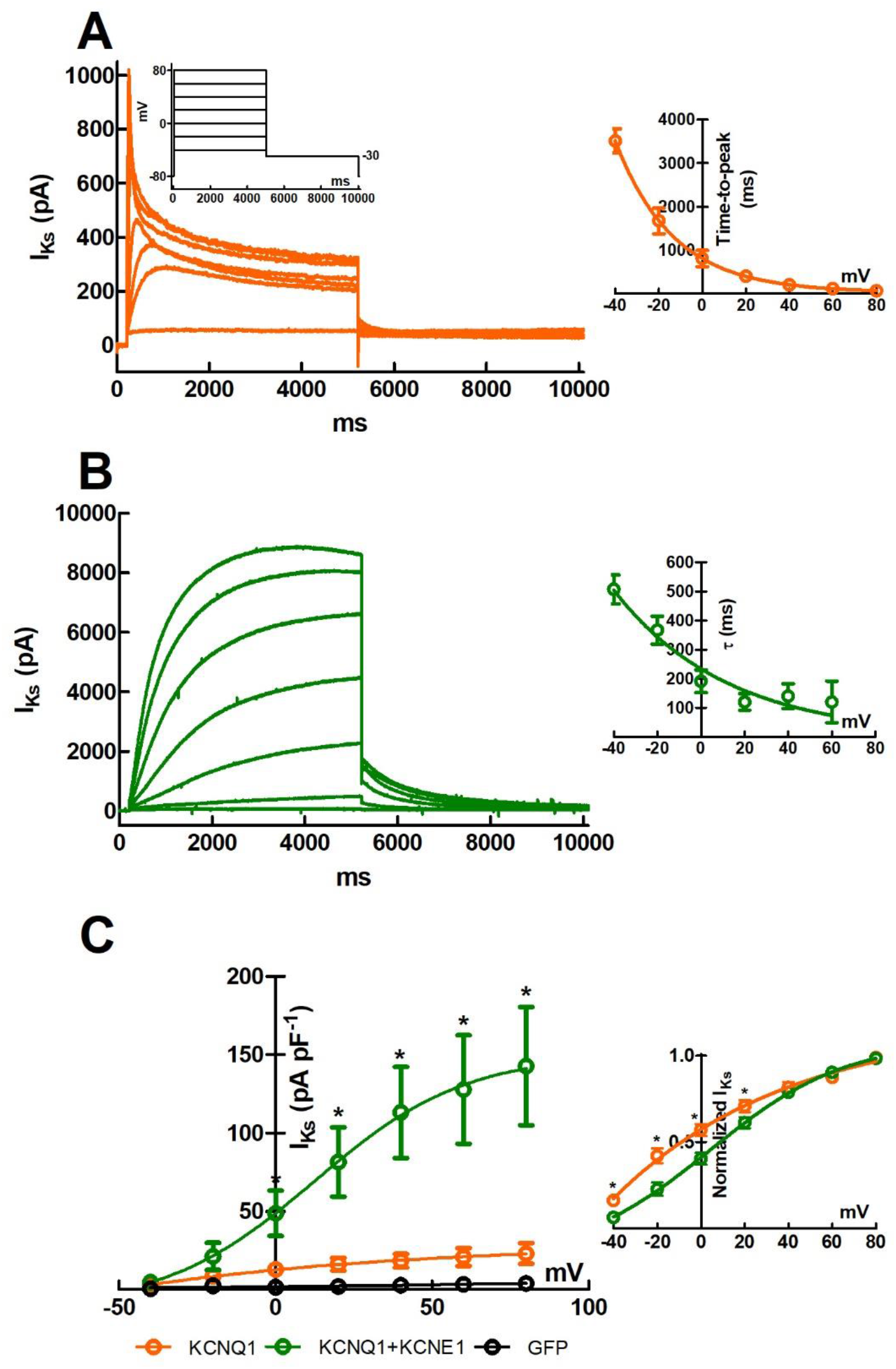
Current-voltage relationship of I_Ks_ generated by zebrafish KCNQ1 and KCNQ+KCNE1 channels in CHO cells. CHO cells were transfected either with KCNQ1 only (1:0) or with KCNQ1 and KCNE1 in 3:1 plasmid ratio. (A) Representative recordings of currents generated by KCNQ1 channels (left) and voltage-dependence of activation kinetics (time-to-peak current) of I_Ks_ (right). (B) Representative recordings of currents generated by KCNQ1+KCNE1 channels (left) and voltage-dependence of activation kinetics (τ) of I_Ks_ (right). (C) Mean (± SEM) current-voltage relationship generated by KCNQ1 and KCNQ1+KCNE1 channels, and cell transfected with GFT plasmid only. The voltage protocol used to elicit the delayed rectifier current is shown in the inset of the panel A. The results are means (± SEM) of 22 and 24 cells for KCNQ1 and KCNQ1+KCNE1, respectively. Asterisks indicate statistically significant differences (*p*<0.05) between the channel assemblies.

### Effects of I_Ks_ inhibitors on homomeric KCNQ1 and heteromeric KCNQ1+KCNE1 channels

Chromanol 293B and its derivative HMR-1556 are blockers of the mammalian slow delayed rectifier K^+^ channels. Chromanol 293B inhibited the zebrafish I_Ks_ current in concentration-dependent manner (**Figure 2A**). The chromanol 293B concentration for half-maximal inhibition (IC_50_) was 13.1 ± 5.8 μM and 13.4 ± 2.8 μM for KCNQ1 and KCNQ1+KCNE1 channels, respectively (*p*>0.05). Complete inhibition was attained at 300 μM chromanol 293B for both channel types. HMR-1556 inhibited I_Ks_ with a much higher affinity than chromanol 293B (**Figure 2B**) with IC_50_-values of 0.1 ± 0.1 μM and 1.5 ± 0.8 μM for KCNQ1 and KCNQ1+KCNE1 channel compositions, respectively. This difference between channel assemblies is statistically significant (*p*=0.01). At the concentration of 100 μM, HMR-1556 completely inhibited both currents.

**FIGURE 2.**
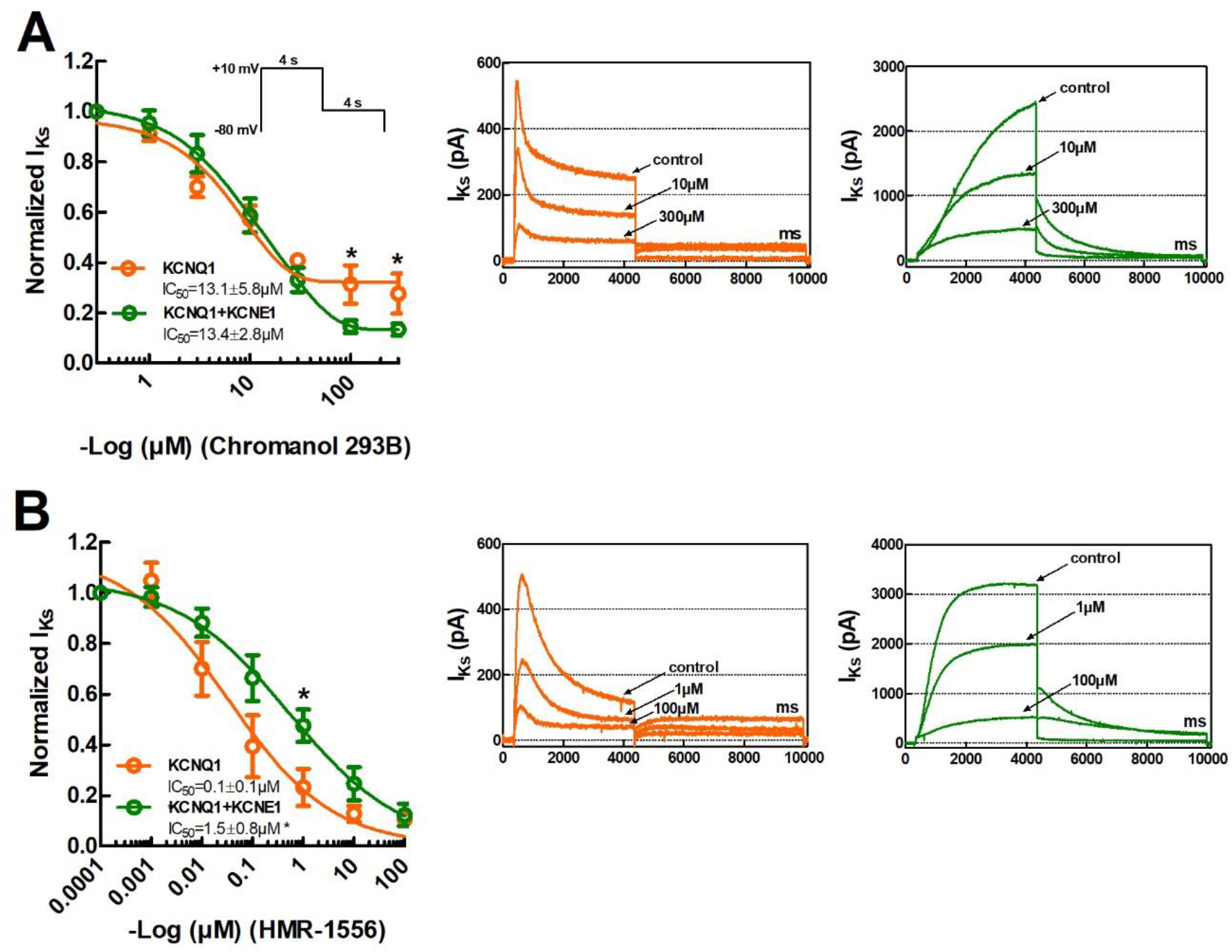
(A) Concentration-response curves of currents generated by zebrafish KCNQ1 and KCNQ+KCNE1 channels in CHO cells to chromanol 293B (left) and representative recordings of I_Ks_ currents in the absence and presence of the drug (right). (B) Concentration-response curves of currents generated by zebrafish KCNQ1 and KCNQ+KCNE1 channels in CHO cells to HMR-1556 (left) and representative recordings of I_Ks_ currents in the absence and presence of the drug (right). The voltage protocol used to elicit the I_Ks_ peak current is shown in the inset of the panel A. Concentration for half-maximal inhibition of the current (IC_50_) is given within the figure. Asterisks indicate statistically differences (*p*<0.05) between the channel assemblies. The results are means (± SEM) of 12-14 cells.

### Effects of IKs activators on homomeric KCNQ1 and heteromeric KCNQ1+KCNE1 channels

R-L3 and mefenamic acid are identified as activators of the mammalian I_Ks_ (Salata et al., 1998; Abitbol et al., 1999; Xu, X. et al., 2002; Wang et al. 2020b). Surprisingly, R-L3 inhibited the zebrafish I_Ks_ at all tested drug concentrations regardless of the type of the channel assembly (**Figure 3A**). R-L3 inhibited I_Ks_ generated by KCNQ1 and KCNQ1+KCNE1 channels with similar affinity (IC_50_ 1.1 ± 0.4 μM and 1.0 ± 0.4 μM, respectively) (*p*>0.05). At the concentration of 10 μM R-L3 almost completely inhibited I_Ks_ of both channel types. At low concentrations (0.001 and 0.01 μM), mefenamic acid slightly stimulated the current generated by the homomeric KCNQ1 channels, while higher concentrations (1-100 μM) inhibited the current (**Figure 3B**). However, mefenamic acid maximally inhibited only about 40% of the current produced by KCNQ1 channels. The current generated by the heteromeric KCNQ1+KCNE1 channels was monotonically and completely inhibited by mefenamic acid (**Figure 3B**). The mefenamic acid concentration for half-maximal inhibition was 9.5 ± 4.8 μM and 3.3 ± 1.8 μM for KCNQ1 and KCNQ1+KCNE1 channels, respectively (*p*>0.05).

**FIGURE 3.**
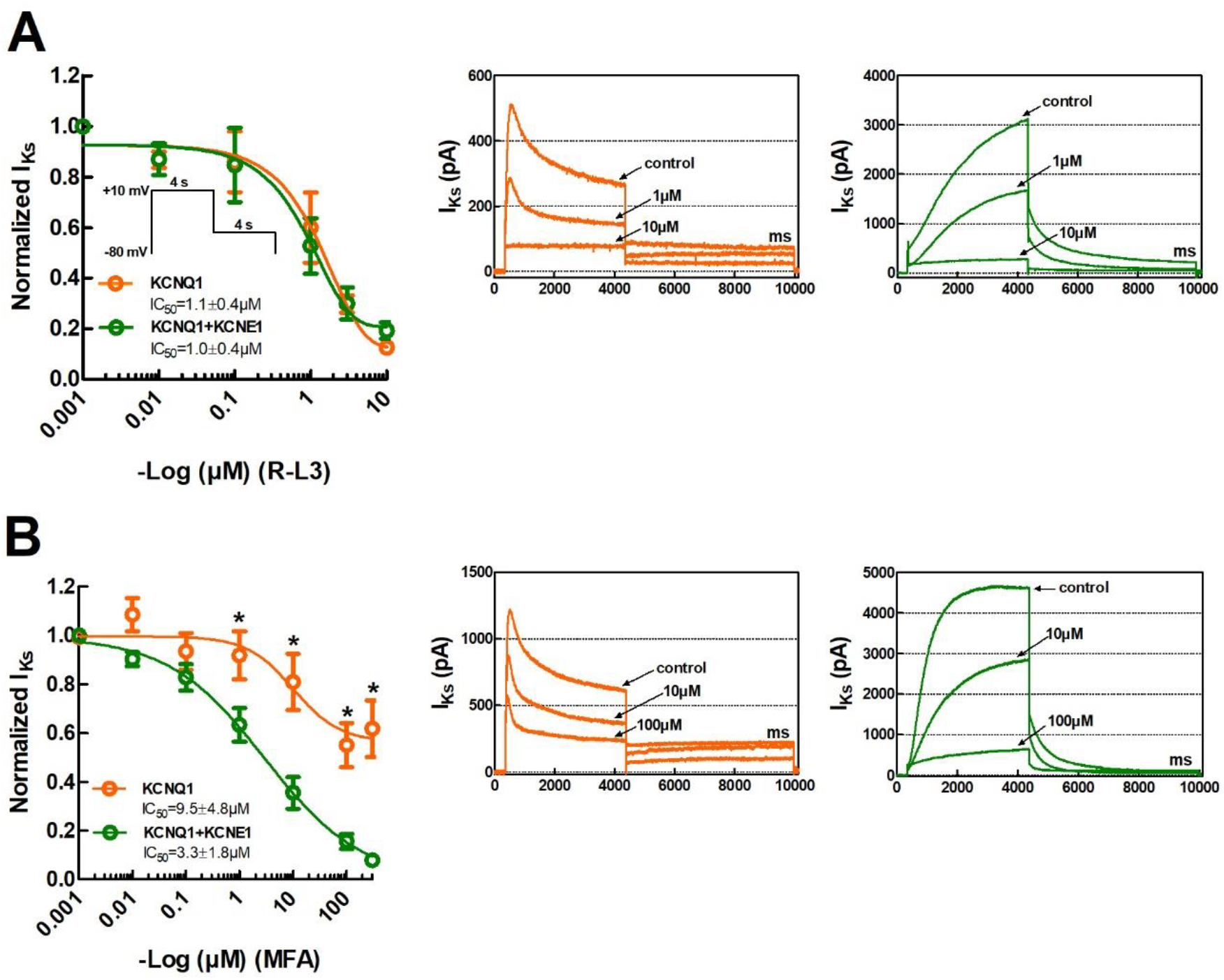
(A) Concentration-response curves of currents generated by zebrafish KCNQ1 and KCNQ+KCNE1 channels in CHO cells to R-L3 (left) and representative recordings of I_Ks_ currents (right). (B) Concentration-response curves of currents generated by zebrafish KCNQ1 and KCNQ+KCNE1 channels in CHO cells to mefenamic acid (left) and representative recordings of I_Ks_ currents (right). The voltage protocol used to elicit the I_Ks_ peak current is shown in the inset of the panel A. Concentration for half-maximal inhibition of the current (IC_50_) is given within the figure. Asterisks indicate statistically differences (*p*<0.05) between the channel assemblies. The results are means (± SEM) of 12-14 cells.

## DISCUSSION

The present results show that inhibitors of I_Ks_, chromanol 293B and HMR-1556, inhibit zebrafish I_Ks_ with a similar potency as they inhibit mammalian I_Ks_ (**Table 1**). In contrast, it was surprising that R-L3 and mefenamic acid, which are identified as activators of mammalian I_Ks_, inhibited the currents generated by the zebrafish KCNQ1 and KCNQ1+KCNE1 channels. Thus, in some respects, the response of the zebrafish I_Ks_ to drugs was similar to that of mammalian I_Ks_, but in some other respects, responses of zebrafish and mammalian I_Ks_ were completely opposite. Also, the channel assembly affected drug responses of zebrafish I_Ks_. Homotetrameric KCNQ1 and KCNQ1+KCNE1 channels were differently by two I_Ks_-modifying drugs, HMR-1556 and mefenamic acid. This is not surprising given that the assembly KCNE1 with KCNQ1 is known to significantly modulate drug responses of I_Ks_ (Wang et al. 2020a).

**Table 1.** Concentrations for half-maximal inhibition (IC_50_) of mammalian and zebrafish I_Ks_ current to chromanol 293B and HMR-1556.

| Species and tissue | Chromanol<br>$IC_{50}$ ($\mu M$) | HMR-1556<br>$IC_{50}$ ($\mu M$) | Reference |
| --- | --- | --- | --- |
| Human KCNQ1 | 26.9 |  | (Lerche et al., 2007) |
| Human KCNQ1/KCNE1 | 6.9 |  | (Lerche et al., 2007) |
| Human KCNQ1 | 65.4 |  | (Bett et al., 2006) |
| Human KCNQ1/KCNE1 | 15.1 |  | (Bett et al., 2006) |
| Guinea-pig ventricular myocytes ($I_{Ks}$) | 2.1 | | (Busch, A. et al., 1996) |
| Guinea-pig KCNQ1/KCNE1 | 6.51 |  | (Printemps et al., 2019) |
| Guinea-pig ventricular myocytes ($I_{Ks}$) | 3.17 | | (Fujisawa et al., 2000) |
| Guinea-pig SA node cells ($I_{Ks}$) | 5.3 | | (Ding et al., 2002) |
| Dog ventricular myocytes ($I_{Ks}$) | 1.8 | | (Sun et al., 2001) |
| Mouse KCNQ1 | 40.9 |  | (Busch et al., 1997) |
| Mouse KCNQ1/KCNE1 | 6.7 |  | (Busch et al., 1997) |
| Dog ventricular myocytes ($I_{Ks}$) | | 0.0105 | (Thomas et al., 2003) |
| Guinea-pig atrial myocytes ($I_{Ks}$) | | 0.061 | (Bosch et al., 2003) |
| Guinea-pig ventricular myocytes ($I_{Ks}$) | | 0.034 | (Gerlach et al., 2001) |
| Zebrafish KCNQ1 | 13.1 | 0.1 | present study |
| Zebrafish KCNQ1/KCNE1 | 13.4 | 1.5 | present study |

### General properties of zebrafish KCNQ1 and KCNQ1+KCNE1 currents

The activation kinetics of the I_Ks_ current and the transcript levels of the KCNQ1 and KCNE1 subunits in the zebrafish ventricle suggest that the I_Ks_ current is probably produced by channel assemblies consisting of a mixture of KCNQ1 homotetramers and KCNQ1+KCNE1 assemblies (3:1) (Abramochkin et al., 2018). Therefore, drug responses of the homotetrameric KCNQ1 channels and the channel assemblies produced by transfection of CHO cells with KCNQ1 and KCNE1 plasmids in 3:1 ratio were examined. The properties of the native ventricular I_Ks_ should be intermediate to those described here for KCNQ1 and KCNQ1+KCNE1 channels in CHO cells. Consistent with the known effects of KCNE1 on the activation kinetics of the I_Ks_ (Barhanin et al., 1996; Sanguinetti et al., 1996), the current generated by homotetrameric KCNQ1 channels was much faster than that generated by KCNQ1+KCNE1 channel assemblies. The KCNQ1 homotetrameric channels produced a prominent early component that inactivated to a steady level during the 5 second depolarizing pulse (Abramochkin et al., 2018). The voltage-dependence of I_Ks_ activation was slightly shifted to more negative voltages, as expected for the heteromeric KCNQ1+KCNE1 channels (Barhanin et al., 1996; Sanguinetti et al., 1996).

### Responses to I_Ks_ blockers

Chromanol 293B and its derivative HMR-1556 are established blockers of the I_Ks_ channels. Indeed, both drugs inhibited the zebrafish I_Ks_ so that approximately 80% of the current was blocked at the concentration of 300 and 100 μM for chromanol 293B and HMR-1556, respectively. The IC_50_-values of the zebrafish channels to chromanol 293B and HMR-1556 were similar as previously reported for the respective mammalian channels. Like the mammalian cardiac I_Ks_, the affinity of HMR-1556 for the zebrafish I_Ks_ channel is almost two orders of magnitude higher than that of chromanol 293B (**Table 1**). The high similarity of chromanol 293B and HMR-1556 affinities towards zebrafish and mammalian I_Ks_ channels is not unexpected considering the high overall sequence similarity of their KCNQ1 proteins (68.3%), the binding target of chromanols. The chromanol receptor lies in the inner pore vestibule of the KCNQ1 channel with binding sites in the pore loop (H5) and the S6 transmembrane domain of the protein. Threonine-312, isoleucine-337 and phenylalanine-340 of the KCNQ1 are the critical bindings sites for chromanol 293B (Lerche et al., 2007). The same amino acid residues exist in the zebrafish KCNQ1 suggesting that the chromanol 293B binding affinity should be similar for zebrafish and human channels (Hassinen et al., 2011; Abramochkin et al., 2018). In fact, the H5 loop and the S6 domain are identical in human and zebrafish KCNQ1 except for the position 324 (isoleucine vs. valine).

Interaction of KCNE1 beta subunit with KCNQ1 alpha subunit is known to increase the inhibition potency of chromanol 293B in mammalian I_Ks_ channels (Busch et al., 1997; Bett et al., 2006; Lerche et al., 2007). For example, in human and mouse, KCNQ1+KCNE1 channels are 4-6 times more sensitive to chromanol 293B than KCNQ1 channels (**Table 1**). It was therefore surprising to find that the chromanol 293B sensitivity of zebrafish I_Ks_ channels was independent of the channel composition; the IC_50_-values of KCNQ1 and KCNQ1+KCNE1 were almost identical. In addition, zebrafish KCNQ1 channels were 15 times more sensitive to HMR-1556 than KCNQ1+KCNE1 channels, as if KCNE1 inhibited HMR-1556 binding to KCNQ1. Because the chromanol binding site of KCNQ1 appears to be identical in mammalian and fish channels, different responses of zebrafish KCNQ1+KCNE1 channels (no enhancement of chromanol 293B block by KCNE1; reduction of HMR-1556 block by KCNE1) compared to the corresponding mammalian channels are likely to be in different effect of KCNE1 on KCNQ1 in the zebrafish channel. This would not be surprising given the small sequence similarity (46.7%) between human and zebrafish KCNE1.

### Responses to I_Ks_ activators

Contrary to expectations, R-L3, an agonists of the mammalian I_Ks_ current (Salata et al., 1998; Xu, X. et al., 2002; Seebohm et al., 2003), inhibited the zebrafish I_Ks_. However, it is important to note that the effect of R-L3 on the mammalian I_Ks_ current is complex. First, R-L3 is a partial agonist and has a biphasic effect on the mammalian I_Ks_. At low concentrations (0.03-1.0 μM) it activates I_Ks_ and at high concentrations (10 μM) it inhibits I_Ks_ (Salata et al., 1998). Second, the effect of R-L3 is stereospecific, the d-enantiomer activates I_Ks_ and the l-enantiomer inhibits I_Ks_ (Salata et al., 1998; Corici et al., 2013). Third, association of KCNQ1 with KCNE1 subunits prevents the activation of channel by R-L3 (Salata et al., 1998). The putative binding site of R-L3 locates in the S5 and S6 transmembrane domains of the KCNQ1 protein (Seebohm et al., 2003). It has been suggested that KCNE1 and R-L3 compete for this binding site which thus explains the effect of channel assembly on I_Ks_. In contrast to the biphasic effect on the mammalian I_Ks_ current, R-L3 had only the inhibitory effect on zebrafish I_Ks_ channels. This effect was independent of the channel structure, as the currents produced by both KCNQ1 and KCNQ1+KCNE1 were reduced with the same drug affinity (IC_50_ about 1 μM). Since S5 and S6 domains of zebrafish and mammalian KCNQ1 are practically identical (Hassinen et al., 2011; Abramochkin et al., 2018), the differences in I_Ks_ responses between mammalian and fish channels are unlikely to be due to the KCNQ1 protein. It is more likely that the zebrafish KCNE1 protein differs so much structurally from its mammalian counterpart that the interaction between KCNQ1 and KCNE1 is different in these vertebrate groups.

Mefenamic acid is an established activator of the mammalian I_Ks_ current at the concentration 100 μM (Busch et al., 1997; Abitbol et al., 1999; Unsöld et al., 2000; Toyoda et al., 2006). Still, at the concentration range of 0.001-300 μM mefenamic acid inhibited the zebrafish I_Ks_, although did not completely abolish it. Mefenamic acid acts extracellularly and causes an easily recognizable change in kinetics and amplitude of I_Ks_. It changes the slowly activating and deactivating I_Ks_ into an almost linear current with instantaneous onset and slowed tail current decay (Abitbol et al., 1999; Toyoda et al., 2006; Wang et al. 2020b). No such changes were found in the zebrafish I_Ks_. In mammalian I_Ks_, mefenamic acid is only effective on heteromeric channels comprising both KCNQ1 and KCNE1 subunits (Busch et al., 1997). Mefenamic acid’s effect on mammalian I_Ks_ requires lysine-41 and a few other surrounding residues on the extra cellular surface of the KCNE1, which might explain why homotetrameric KCNQ1 channels are insensitive to this drug (Abitbol et al., 1999). Indeed, three residues lysine-41, leucine-42 and glutamic acid-43 seem to form the critical sequence in the mammalian KCNE1 for activation by mefenamic acid. Notably, in the zebrafish KCNE the residues in positions 41-43 are histidine, leucine and serine, respectively (Hassinen et al., 2011; Abramochkin et al., 2018). Therefore, it is likely that the interaction of KCNE1 with KCNQ1 is different from that of the mammalian I_Ks_ channel. Expression of the zebrafish KCNQ1 together with the mammalian KCNE1 would probably reveal the role zebrafish KCNE1 in the response of I_Ks_ to mefenamic acid and possibly to R-L3.

### Implications for the use of zebrafish as a preclinical drug screening model

Cardiac AP is generated by the delicate interaction of several inward and outward ion currents of the sarcolemma. Under normal unstressed conditions, the role of I_Ks_ is probably minor in determining the shape of zebrafish cardiac AP (Abramochkin et al., 2018). Therefore, one might think that the atypical responses of zebrafish I_Ks_ to drugs (exemplified by responses to R-L3 and mefenamic acid) do not necessarily indicate any severe limitation for drug screening. Because the atypical drug responses of zebrafish I_Ks_ may be due in part to a significantly different sequence of the KCNE1 subunit, particularly its interacting binding domain with KCNQ1, this problem could be easily eliminated by creating a transgenic zebrafish. The problem in this scenario is that the I_Ks_ channel is not likely to be the only ion channel that should be altered (Verkerk and Remme, 2012; Hassinen et al., 2015). Given the stringent criteria for drug screening (Wall and Shani, 2008), zebrafish may not be an optimal and safe general-purpose model for screening drug molecules for adult humans. The ion channel composition (e.g. large T-type Ca^2+^ current), myocyte structure (absence of T-tubuli) and management of intracellular free Ca^2+^ concentration (less dependent on sarcoplasmic reticulum Ca^2+^ release) of the zebrafish heart is much more similar to neonatal than adult mammalian heart (Brette et al., 2008; Nemtsas et al., 2010; Verkerk and Remme, 2012; Bovo et al., 2013; Hassinen et al., 2015). Therefore, zebrafish might be a useful translational model for cardiac electrophysiology of fetal and neonatal individuals and possibly in addressing drug effects on fetal and neonatal human hearts (Vornanen et al., 2018).

### Limitations of the study

This is the first study to look at the effects of drugs on the zebrafish I_Ks_ and highlights some research topics that were not addressed in this study. Although the cardiac I_Ks_ is considered to be generated by channels that constitute of KCNQ1 and KCNE1 subunits, it is possible that that other KCNE subunits (KCNE2-5) will affect the phenotype of I_Ks_ current (Jespersen et al., 2005; Roura-Ferrer et al., 2010). Therefore, drug effects should be examined on APs of zebrafish heart in the presence of I_Kr_ blockers and under beta-adrenergic activation. Generating chimeras of zebrafish KCNQ1 and human KCNE1 or mutating the putative KCNQ1 binding domain in zebrafish KCNE1 could reveal the significance of zebrafish KCNE1 sequence in drug responses.

## Acknowledgements

Anita Kervinen is acknowledged for her excellent technical assistance.

## Author contributions

Conceptualization: M.V.; Methodology: J.H., M.H., M.V.; Investigation: J.H., M.H.; Writing – original draft: M.V.; Writing – review & editing: J.H., M.H., M.V.; Project administration: M.V.; Funding acquisition: M.V.

## References

Abitbol, I., Peretz, A., Lerche, C., Busch, A. E. and Attali, B. (1999). Stilbenes and fenamates rescue the loss of IKS channel function induced by an LQT5 mutation and other IsK mutants. EMBO J. 18, 4137–4148

Abramochkin, D. V., Hassinen, M. and Vornanen, M. (2018). Transcripts of Kv7.1 and MinK channels and slow delayed rectifier K+ current (I_Ks_) are expressed in zebrafish *(Danio rerio)* heart. Pflüg. Arch. 470, 1753–1764

Barhanin, J., Lesage, F., Guillemare, E., Fink, M., Lazdunski, M. and Romey, G. (1996). KvLQT1 and IsK (minK) proteins associate to form the I_Ks_ cardiac potassium current. Nature 384, 78–80

Bendahhou, S., Marionneau, C., Haurogne, K., Larroque, M. M., Derand, R., Szuts, V., Escande, D., Demolombe, S. and Barhanin, J. (2005). In vitro molecular interactions and distribution of KCNE family with KCNQ1 in the human heart. Cardiovascular Research 67, 529–538

Bett, G. C. L., Morales, M. J., Beahm, D. L., Duffey, M. E. and Rasmusson, R. L. (2006). Ancillary subunits and stimulation frequency determine the potency of chromanol 293B block of the KCNQ1 potassium channel. J. Physiol. 576, 755–767

Bett, G. C. L. and Rasmusson, R. L. (2008). Modification of K channel–drug interactions by ancillary subunits. J. Physiol. 586, 929–950

Bosch, R. F., Schneck, A. C., Csillag, S., Eigenberger, B., Gerlach, U., Brendel, J., Lang, H. J., Mewis, C., Gögelein, H. and Seipel, L. (2003). Effects of the chromanol HMR 1556 on potassium currents in atrial myocytes. Naunyn Schmiedebergs Arch. Pharmacol. 367, 281–288

Bovo, E., Dvornikov, A. V., Mazurek, S. R., de Tombe, P. P. and Zima, A. V. (2013). Mechanisms of Ca2+ handling in zebrafish ventricular myocytes. Pflügers Arch. 465, 1775–1784

Brette, F., Luxan, G., Cros, C., Dixey, H., Wilson, C. and Shiels, H. A. (2008). Characterization of isolated ventricular myocytes from adult zebrafish *(Danio rerio)*. Biochem. Biophys. Res. Commun. 374, 143–146

Busch, A., Suessbrich, H., Waldegger, S., Sailer, E., Greger, R., Lang, H., Lang, F., Gibson, K. and Maylie, J. (1996). Inhibition of I_Ks_ in guinea pig cardiac myocytes and guinea pig I_sK_ channels by the chromanol 293B. Pflügers Archiv 432, 1094–1096

Busch, A. E., Busch, G. L., Ford, E., Suessbrich, H., Lang, H. J., Greger, R., Kunzelmann, K., Attali, B. and Sthhmer, W. (1997). The role of the IsK protein in the specific pharmacological properties of the I_Ks_ channel complex. Br. J. Pharmacol. 122, 187–189

Chen, H., Kim, L. A., Rajan, S., Xu, S. and Goldstein, S. A. N. (2003). Charybdotoxin binding in the I_Ks_ pore demonstrates two MinK subunits in each channel complex. Neuron 40, 15–23

Corici, C., Kohajda, Z., Kristóf, A., Horváth, A., Virág, L., Szél, T., Nagy, N., Szakonyi, Z., Fülöp, F. and Muntean, D. M. (2013). L-364,373 (R-L3) enantiomers have opposite modulating effects on I Ks in mammalian ventricular myocytes. Can. J. Physiol. Pharmacol. 91, 586–592

de la Cruz, A., Perez-Rodriguez, M. E., Rainer, Q. C., Liin, S. I. and Larsson, P. H. (2020). Zebrafish Heart as a Model for Early-Screening of Human Antiarrhythmic Drugs. Biophys. J. 118, 114a

Ding, W. G., Toyoda, F. and Matsuura, H. (2002). Blocking action of chromanol 293B on the slow component of delayed rectifier K current in guinea-pig sino-atrial node cells. Br. J. Pharmacol. 137, 253–262

Fujisawa, S., Ono, K. and Iijima, T. (2000). Time-dependent block of the slowly activating delayed rectifier K current by chromanol 293B in guinea-pig ventricular cells. Br. J. Pharmacol. 129, 1007–1013

Gerlach, U., Brendel, J., Lang, H. J., Paulus, E. F., Weidmann, K., Brüggemann, A., Busch, A. E., Suessbrich, H., Bleich, M. and Greger, R. (2001). Synthesis and activity of novel and selective IKs-channel blockers. J. Med. Chem. 44, 3831–3837

Hassinen, M., Haverinen, J., Hardy, M. E., Shiels, H. A. and Vornanen, M. (2015). Inward rectifier potassium current (I_K1_) and Kir2 composition of the zebrafish *(Danio rerio)* heart. Pflüg. Arch. 467, 2437–2446

Hassinen, M., Laulaja, S., Paajanen, V., Haverinen, J. and Vornanen, M. (2011). Thermal adaptation of the crucian carp *(Carassius carassius)* cardiac delayed rectifier current, I_Ks_, by homomeric assembly of Kv7.1 subunits without MinK. Am. J. Physiol. 301, R255–R2665

Honek, J. (2017). Preclinical research in drug development. Med. Writing 26, 5–8

Jespersen, T., Grunnet, M. and Olesen, S. P. (2005). The KCNQ1 Potassium Channel: From Gene to Physiological Function. Physiology 20, 408–416

Jost, N., Virag, L., Bitay, M., Takacs, J., Lengyel, C., Biliczki, P., Nagy, Z., Bogats, G., Lathrop, D. A., Papp, J. G. et al. (2005). Restricting excessive cardiac action potential and QT prolongation: A vital role for I_Ks_ in human ventricular muscle. Circulation 112, 1392–1399

Joukar, S. (2021). A comparative review on heart ion channels, action potentials and electrocardiogram in rodents and human: extrapolation of experimental insights to clinic. Lab. Anim. Res. 37, 1–15

Kari, G., Rodeck, U. and Dicker, A. P. (2007). Zebrafish: an emerging model system for human disease and drug discovery. Clin. Pharmcol. Ther. 82, 70–80

Lerche, C., Bruhova, I., Lerche, H., Steinmeyer, K., Wei, A. D., Strutz-Seebohm, N., Lang, F., Busch, A. E., Zhorov, B. S. and Seebohm, G. (2007). Chromanol 293B binding in KCNQ1 (Kv7.1) channels involves electrostatic interactions with a potassium ion in the selectivity filter. Mol. Pharmacol. 71, 1503–1511

MacRae, C. A. and Peterson, R. T. (2015). Zebrafish as tools for drug discovery. Nature Reviews Drug Discovery 14, 721–731

Morin, T. J. and Kobertz, W. R. (2008). Counting membrane-embedded KCNE beta-subunits in functioning K+ channel complexes. Proc. Natl. Acad. Sci. U. S. A. 105, 1478–1482

Narumanchi, S., Wang, H., Perttunen, S., Tikkanen, I., Lakkisto, P. and Paavola, J. (2021). Zebrafish Heart Failure Models. Front. Cell Dev. Biol. 9, 1061

Nemtsas, P., Wettwer, E., Christ, T., Weidinger, G. and Ravens, U. (2010). Adult zebrafish heart as a model for human heart? An electrophysiological study. J. Mol. Cell. Cardiol. 48, 161–171

Parng, C., Seng, W. L., Semino, C. and McGrath, P. (2002). Zebrafish: a preclinical model for drug screening. Assay Drug. Dev. Techn. 1, 41–48

Pound, P. and Ritskes-Hoitinga, M. (2018). Is it possible to overcome issues of external validity in preclinical animal research? Why most animal models are bound to fail. J. Transl. Med. 16, 1–8

Printemps, R., Salvetat, C., Faivre, J. F., Le Grand, M., Bois, P. and ou Maati, H. M. (2019). Role of Cardiac I_Ks_ Current in Repolarization Reserve Process During Late Sodium Current (I_NaL_) Activation. Cardiol. Cardiov. Med. 3, 168–185

Rocchetti, M., Besana, A., Gurrola, G. B., Possani, L. D. and Zaza, A. (2001). Rate dependency of delayed rectifier currents during the guinea-pig ventricular action potential. J. Physiol. 534, 721–732

Roura-Ferrer, M., Solé, L., Oliveras, A., Dahan, R., Bielanska, J., Villarroel, Á, Comes, N. and Felipe, A. (2010). Impact of KCNE subunits on KCNQ1 (Kv7. 1) channel membrane surface targeting. J. Cell. Physiol. 225, 692–700

Salata, J. J., Jurkiewicz, N. K., Wang, J., Evans, B. E., Orme, H. T. and Sanguinetti, M. C. (1998). A novel benzodiazepine that activates cardiac slow delayed rectifier K+ currents. Mol. Pharmacol. 54, 220–230

Sanguinetti, M. C., Curran, M. E., Zou, A., Shen, J., Specter, P. S., Atkinson, D. L. and Keating, M. T. (1996). Coassembly of KVLQT1 and minK (IsK) proteins to form cardiac I_Ks_ potassium channel. Nature 384, 80–83

Seebohm, G., Pusch, M., Chen, J. and Sanguinetti, M. C. (2003). Pharmacological Activation of Normal and Arrhythmia-Associated Mutant KCNQ1 Potassium Channels. Circulation Research 93, 941–947

Sun, Z. Q., Thomas, G. P. and Antzelevich, C. (2001). Chromanol 293B inhibits slowly activating delayed rectifier and transient outward currents in canine left ventricular myocytes. J. Cardiovasc. Electrophysiol. 12, 472–478

Thomas, G. P., Gerlach, U. and Antzelevitch, C. (2003). HMR 1556, a potent and selective blocker of slowly activating delayed rectifier potassium current. J. Cardiovasc. Pharmacol. 41, 140–147

Toyoda, F., Ueyama, H., Ding, W. G. and Matsuura, H. (2006). Modulation of functional properties of KCNQ1 channel by association of KCNE1 and KCNE2. Biochem. Biophys. Res. Commun. 344, 814–820

Unsöld, B., Kerst, G., Brousos, H., Hübner, M., Schreiber, R., Nitschke, R., Greger, R. and Bleich, M. (2000). KCNE1 reverses the response of the human K+ channel KCNQ1 to cytosolic pH changes and alters its pharmacology and sensitivity to temperature. Pflügers Archiv 441, 368–378

Verkerk, A. O. and Remme, C. A. (2012). Zebrafish: a novel research tool for cardiac (patho)electrophysiology and ion channel disorders. Front. Physiol. 3, 255

Vornanen, M. and Hassinen, M. (2016). Zebrafish heart as a model for human cardiac electrophysiology. Channels 10, 101–110

Vornanen, M., Haverinen, J. and Hassinen, M. (2018). Excitation and excitation-contraction coupling of the zebrafish heart: implications for the zebrafish model in drug screening. In Recent Advances in Zebrafish Researches (ed. Y. Bozkurt), pp. 83–100

Wall, R. J. and Shani, M. (2008). Are animal models as good as we think? Theriogenology 69, 2–9

Wang, K., Terrenoire, C., Sampson, K. J., Iyer, V., Osteen, J. D., Lu, J., Keller, G., Kotton, D. N. and Kass, R. S. (2011). Biophysical properties of slow potassium channels in human embryonic stem cell derived cardiomyocytes implicate subunit stoichiometry. J. Physiol. 589, 6093–6104

Wang, Y., Eldstrom, J. and Fedida, D. (2020a). Gating and regulation of KCNQ1 and KCNQ1 KCNE1 channel complexes. Front. Physiol. 11, 504

Wang, Y., Eldstrom, J. and Fedida, D. (2020b). The I_Ks_ Ion Channel Activator Mefenamic Acid Requires KCNE1 and Modulates Channel Gating in a Subunit-Dependent Manner. Mol. Pharmacol. 97, 132–144

Xu, H., Guo, W. and Nerbonne, J. M. (1999). Four kinetically distinct depolarization-activated K+currents in adult mouse ventricular myocytes. J. Gen. Physiol. 113, 661–677

Xu, X., Salata, J. J., Wang, J., Wu, Y., Yan, G. X., Liu, T., Marinchak, R. A. and Kowey, P. R. (2002). Increasing I Ks corrects abnormal repolarization in rabbit models of acquired LQT2 and ventricular hypertrophy. Am. J. Physiol. 283, H664–H670

